# Multimodal imaging links molecular identity with nanoscale-resolved 3D cell morphology in opossum cortical cultures

**DOI:** 10.64898/2026.11.06.764880

**Authors:** Zahra Ayar, Marta Pongrac, Miranda Mladinic, Jelena Ban, Georg E. Fantner

**Affiliations:** Institute of Bioengineering, School of Engineering, Swiss Federal Institute of Technology Lausanne (EPFL), Lausanne 1015, Switzerland; Faculty of Biotechnology and Drug Development, University of Rijeka, Radmile Matejčić 2, 51000 Rijeka, Croatia

**Keywords:** Scanning ion conductance microscopy, morphology, neurite regeneration, morphometric characterization, multimodal imaging

## Abstract

In the mammalian central nervous system (CNS), the ability to regenerate after injury drops sharply in the first weeks after birth, but how this change is reflected in cell morphology and organization is still not fully understood. Here we cultivated cortical cell cultures from the gray short-tailed opossum (*Monodelphis domestica*) and compared cells derived from early postnatal pups (postnatal day 3-6, “P*_young_*”) with those from older pups (postnatal day 16-18, “P*_old_*”), which differ in their capacity to regenerate after injury. For each culture, we assessed marker-defined cellular composition using fluorescently labeled markers for neurons, glia, and immature/progenitor cells, and characterized their nanoscale-resolved three-dimensional morphology using label-free scanning ion conductance microscopy (SICM). P*_young_* cultures exhibited active change in shape over time, contained a higher proportion of immature SOX2⁺ cells, and grew more neurites (TUJ1⁺) into the lesion site after injury. Moreover, P*_old_* cultures showed more stable soma size (e.g., thickness, area, volume), a larger astroglial (SOX9⁺) population early in culture, and a glia-dominated lesion response. These early astroglia included more immature or reactive (vimentin⁺) astrocytes. SICM further revealed cellular bridges across the lesion, ridges left in the substrate by the injury, and aligned neurite outgrowth. Across all measurements, changes in molecular markers were associated with corresponding changes in cell morphology. This multimodal combination of fluorescence labeling and SICM therefore relates marker-defined cellular composition to physical organization across micro- and nanoscale features during and after postnatal *in vitro* development.

**Graphical Abstract:** 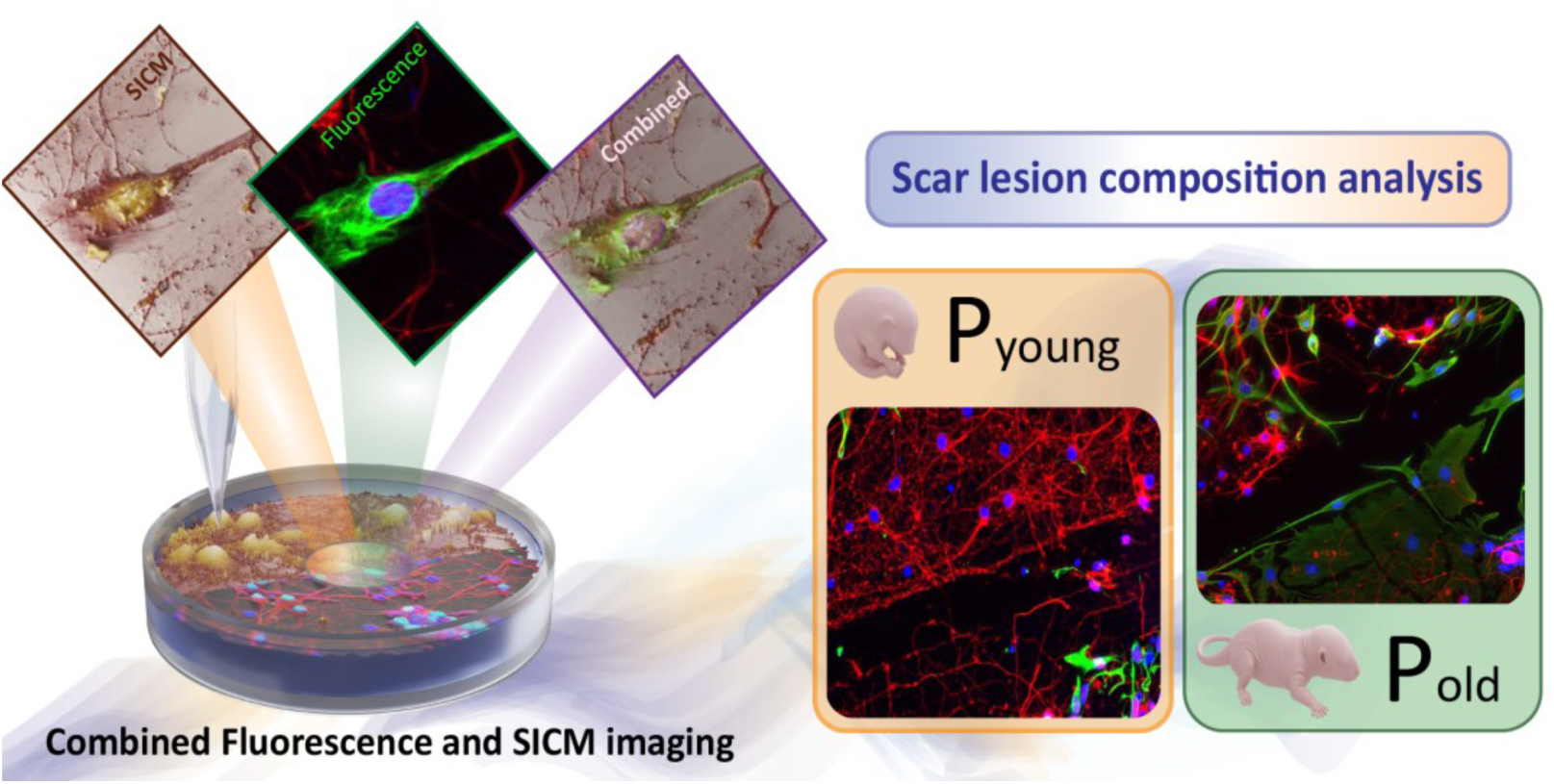

## Introduction

In the adult mammalian central nervous system (CNS), axons generally fail to regrow after injury, which leads to permanent loss of function^1–4^. Regeneration requires reactivating developmental processes that are suppressed as the CNS matures^2,3,5^. Extensive work has identified molecular pathways that regulate intrinsic axon growth and extrinsic inhibition^1,2,6–8^. Neurite regeneration also requires coordinated three-dimensional remodeling, including growth cone expansion^9,10^, cytoskeletal rearrangement^10^, and navigation through a changing environment^5,11^. Nevertheless, how developmental molecular identity relates to three-dimensional shape remains poorly understood.

During CNS development, neurons and glia profoundly reorganize their structure^12^. Axon growth is led by the growth cone, a temporary structure that forms during development and disappears once synapses are made. The ability to form a new growth cone after injury is key to successful regeneration^9,10,13,14^, whereas the reactive glia build-up contributes to scar formation and limits repair^4,11,12,15^. Regenerative capacity therefore depends on gene expression and marker-defined cell identity, but also on the physical organization of neurons, glia, and the injury site.

Fluorescence microscopy and immunocytochemistry are the standard tools for identifying cell types and injury-associated molecular states^16^. However, they rely on labeling specific molecular targets, providing a partial view of the cellular structures. Moreover, conventional optical imaging is limited by diffraction and by poor axial resolution, so it cannot fully capture fine three-dimensional structure^17,18^. For example, common astrocyte markers such as glial fibrillary acidic protein (GFAP) label around 15% of the cell volume and can underrepresent its true volume^19,20^. A method that resolves cell shape directly, without labels, would therefore complement fluorescence-based analysis of cell cultures.

Label-free scanning probe methods can image cell structure directly at high resolution. Scanning ion conductance microscopy (SICM) is one such method that maps the cell surface in three dimensions under physiological conditions without physical contact between the probe and the sample. SICM uses an electrolyte-filled nanopipette and detects the drop in ion current as the nanopore approaches the surface, producing non-contact topographic images with sub-5 nm resolution along the vertical axis^21–23^. Because it does not touch the cell, SICM can image delicate membrane structures and follow live-cell dynamics without deforming them^23–27^. In neurons, SICM has resolved micro- and nanoscale structural dynamics in hippocampal cells^26^, redirected growth cones^28^, measured mechanical properties^29^, and delivered local mechanical stimulation to sensory neurons^30^.

Here, we combine SICM with fluorescence microscopy to link marker-defined cellular composition with three-dimensional cell morphology and surface topography at micro- and nanoscale resolution. Fluorescence labeling enabled the identification of distinct cell subpopulations within the culture, including neurons, glia, and immature or progenitor cells, while SICM measures their morphology without labels. Together, these methods let us link cellular composition to structural changes during *in vitro* development and after injury.

We used the gray short-tailed opossum (*Monodelphis domestica*) as a model organism. Opossums are born immature, at a stage comparable to an early rodent embryo, and complete much of their CNS development after birth^31^. This provides a long postnatal window to study regenerative capacity directly^31–33^. Early postnatal animals regenerate CNS axons well, while older animals shift to a non-regenerative state^34–36^. In the spinal cord, axon regeneration persists through about the first two postnatal weeks^37,38^, compared with only a few days in rodents^39–42^. The opossum also offers practical advantages. Pups develop externally, so their exact developmental age is known, each pregnancy produces many pups, and the mother is not sacrificed, all of which support the 3R principles^43,44^. Because its cortical cells can be grown in dissociated culture^45^, the opossum lets us compare early and later postnatal stages of the same species side by side.

Because regeneration reactivates growth-associated processes that occur during development^46–48^, we reasoned that developmental stage should be visible not only in molecular marker profiles, but also in the three-dimensional shape of neurons, glia, and the lesion environment. We therefore combined fluorescently labeled markers with label-free SICM to compare early and later postnatal opossum cortical cultures, both as they developed *in vitro* and after a mechanical lesion. This multimodal strategy let us relate marker profiles to soma shape, membrane surface, neurite structure, and lesion-site organization at the micro- and nanoscale, and examine whether injury responses linked to regeneration are accompanied by measurable differences in cell shape.

## 2. Results

### 2.1 Study design

We cultivated primary cortical cultures from young (P3-6, “P*_young_*”) and older (P16-18, “P*_old_*”) *Monodelphis domestica* pups under identical conditions and studied them at set days *in vitro* (DIV), covering early, intermediate, mid-to-late, and late stages of *in vitro* development in culture (Figure 1.a; Figure S1). At selected stages, we used fluorescence microscopy to quantify neurons, progenitor-associated cells, and glia, and SICM to record label-free three-dimensional height maps of cell and lesion-site shape (Figure 1.b). Sample sizes for each measurement are given in the corresponding sections and in the Methods section. This design lets us compare molecular profile, three-dimensional shape, and injury response between P*_young_* and P*_old_* cultures.

**Figure 1.**
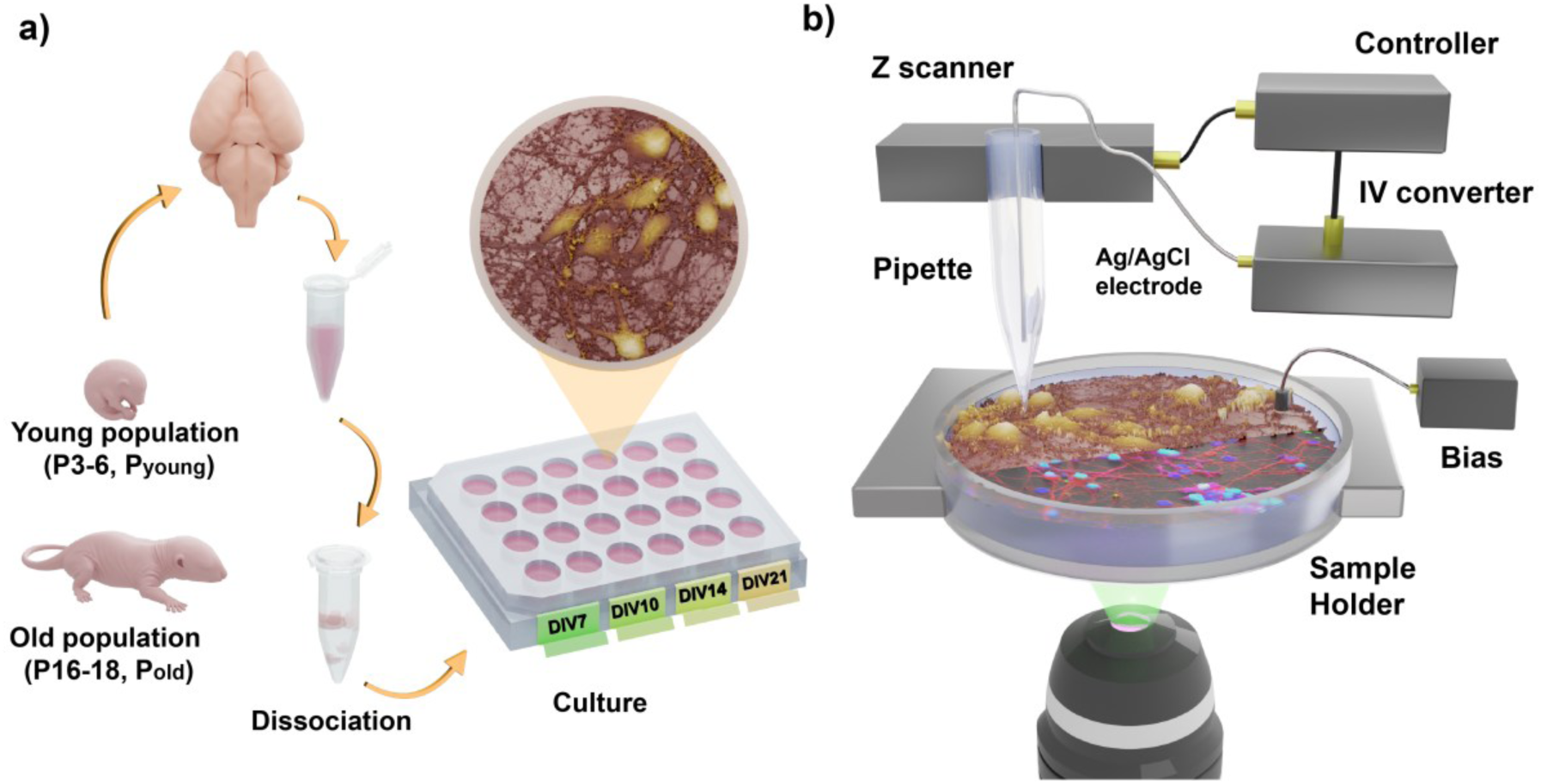
Experimental workflow and imaging strategy. a) Cortical neurons isolated from young (P*_young_*) and older (P*_old_*) *Monodelphis domestica* pups and grown to defined stages. b) SICM setup used for label-free topographic imaging together with fluorescence microscopy.

### 2.2 SICM captures cell structure across *in vitro* development

We used SICM to follow three-dimensional surface structure as the cultures developed. In P*_young_* cultures imaged from DIV1 to DIV21, organization changed progressively (Figure 2.a; Figure S2). At DIV1, cells were mostly isolated and morphologically simple, with limited neurite growth. At later stages, the cultures showed larger somata, denser neurites, and more elaborate neuronal networks. High-resolution SICM resolved submicrometric fine surface features, including neurite extensions, filopodia-like protrusions, and neuron to glia network structures, in both P*_young_* and P*_old_* cultures (Figure 2.b). SICM can therefore capture label-free three-dimensional features throughout *in vitro* cortical culture development.

**Figure 2.**
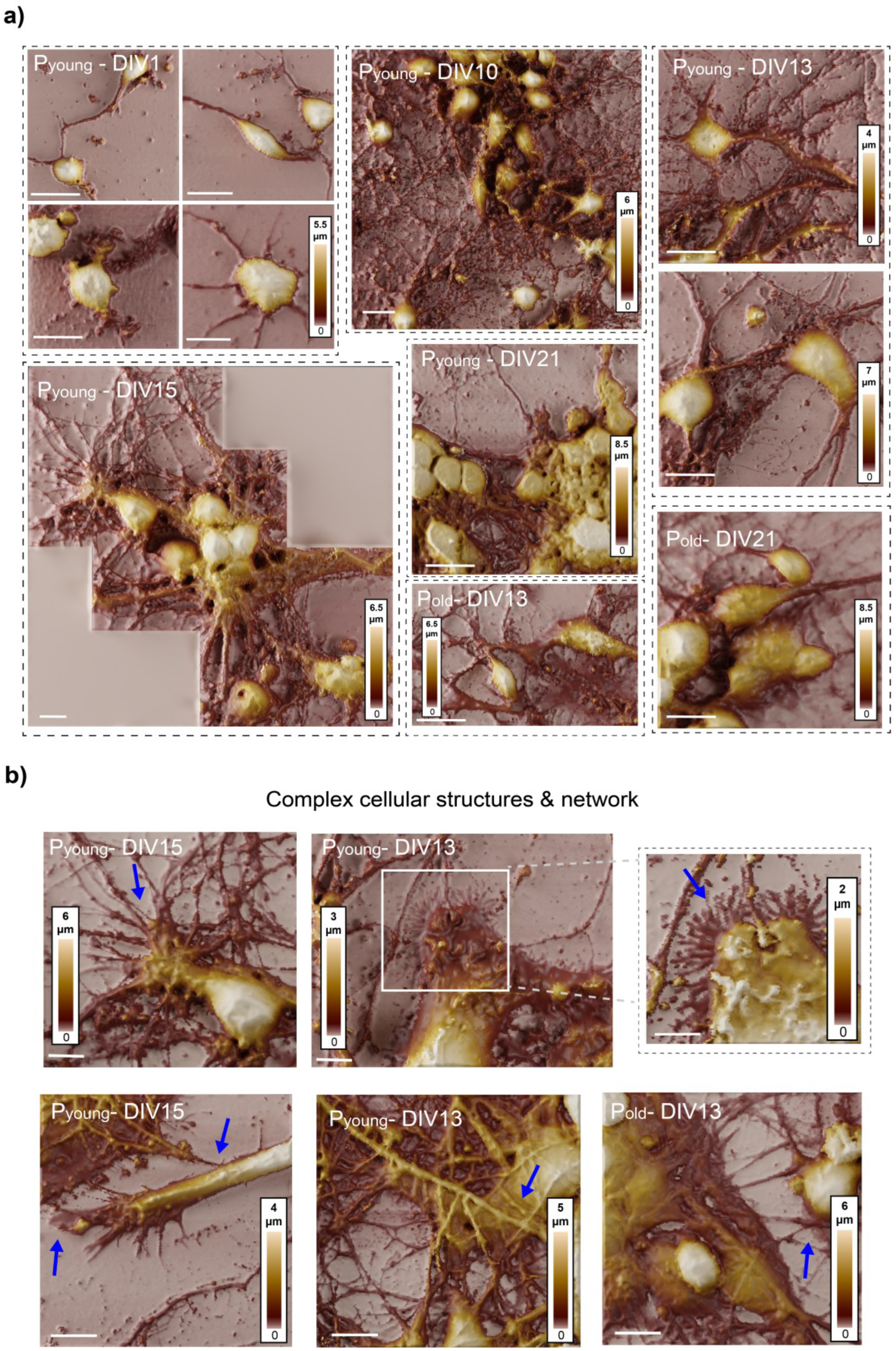
SICM images of cortical cultures across *in vitro* development. a) SICM images of P*_young_* cultures at DIV1, DIV10, DIV13, DIV15, and DIV21, and P*_old_* cultures at DIV13 and DIV21 showing progressive elaboration of neuronal networks with *in vitro* development. b) Representative high-resolution SICM images acquired at intermediate- to-late developmental stages in P*_young_* and P*_old_* mixed neuronal-glial cultures, highlighting fine cellular surface features, interconnected cellular processes, and complex network-like architectures. Blue arrows indicate selected surface protrusions and neurite-like extensions. Scale bar, 10 μm for all images.

### 2.3 Age- and stage-dependent cellular composition and soma morphology

We characterized the neuronal and progenitor composition of the cultures using three markers (Figure 3.a). SRY-box transcription factor 2 (SOX2) is expressed in neural progenitor and immature cells and decreases as cells differentiate ^49–51^. βIII-tubulin (TUJ1) was used as a marker of developing neurons^52^, whereas vimentin was used to identify immature and reactive glial cells, with its expression increasing following CNS injury^53^. We selected these markers based on previous *ex vivo* work in opossum, which showed that progenitor and neuronal markers are co-expressed at distinct stages of postnatal CNS development^54^. We then quantified SOX2⁺ cells co-expressing TUJ1.

**Figure 3.**
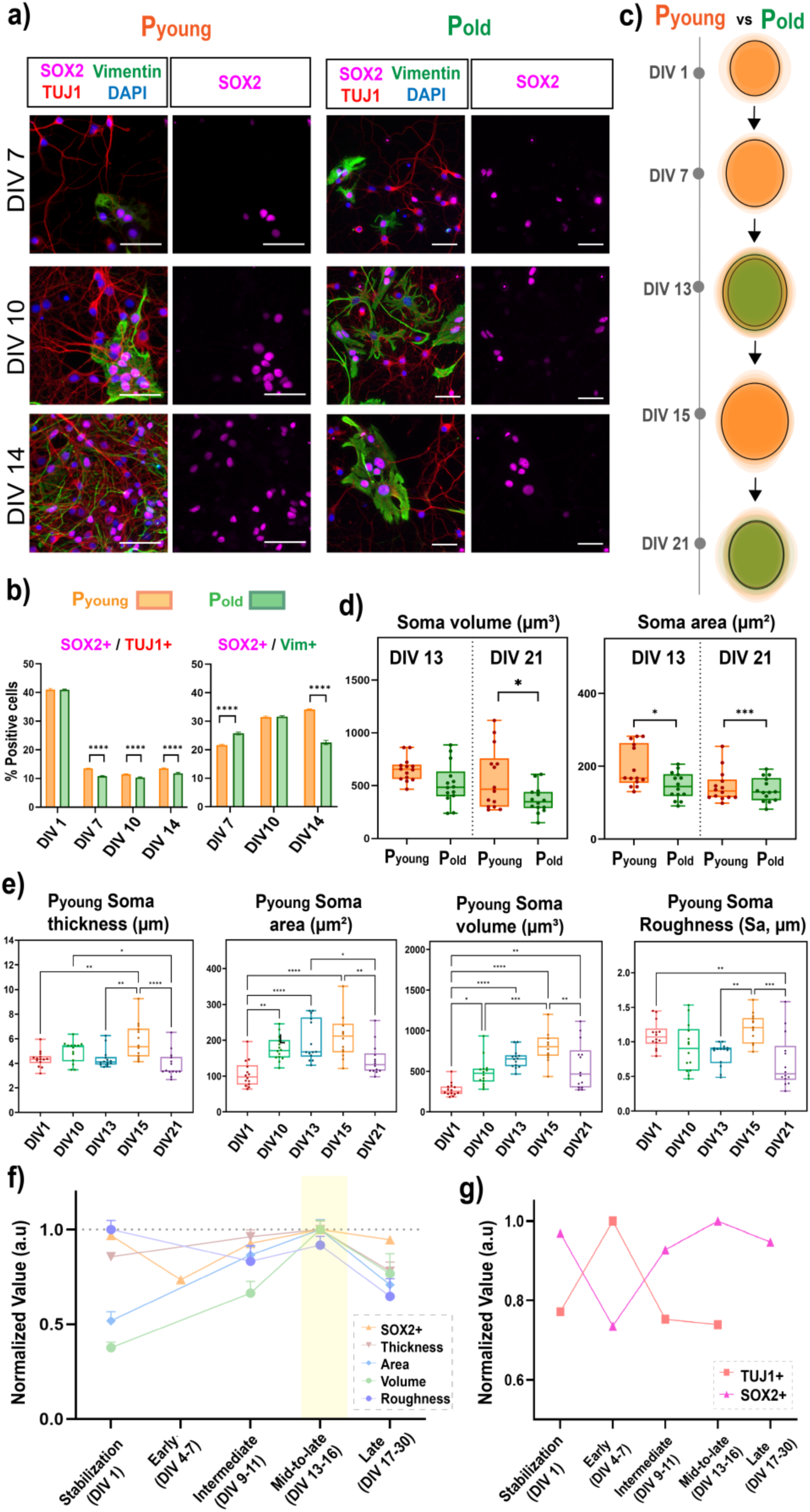
Cellular composition and soma size in P*_young_* and P*_old_* cultures across *in vitro* development. a) Fluorescence microscopy images of P*_young_* and P*_old_* cultures at DIV7, DIV10, and DIV14: SOX2 (magenta), TUJ1 (red), vimentin (green), DAPI (blue). Scale bar, 50 µm, b) SOX2⁺/TUJ1⁺ (left) and SOX2⁺/vimentin⁺ (right) subpopulations across development. SOX2⁺/TUJ1⁺ subpopulation declined sharply after DIV1, whereas the SOX2⁺/vimentin⁺ subpopulation increased in P*_young_* over time. Two-way ANOVA showed significant effects of stage and age group, with significant P*_young_* - P*_old_* differences at most matched stages; for clarity, only the selected comparisons are marked (p < 0.0001). c) Schematic of soma size progression in P*_young_* and P*_old_* cultures, showing proportional growth and partial reduction at later developmental stages. d) Soma area and volume in P*_young_* and P*_old_* cultures at DIV13 and DIV21. At DIV13, P*_young_* somata had a larger projected area than P*_old_* somata (p < 0.05). At DIV21, soma volume was significantly larger in P*_young_* cultures (p < 0.05; n = 14 cells per group). e) Morphometric parameters from SICM height maps of P*_young_* cultures (one-way ANOVA, p < 0.001; n=14 cells per group). Thickness area, volume, and roughness reached a peak value at DIV15, followed by a decline at DIV21. (Tukey post-hoc; *p < 0.05, **p < 0.01, ***p < 0.001, ****p < 0.0001). f) Normalized SICM measured parameters and the proportion of SOX2⁺ cells across *in vitro* development. The SICM parameters followed similar developmental trends and reached their highest normalized values around DIV15, coinciding with the maximum proportion of SOX2⁺ cells. The measurements converged during the mid- to late developmental stages (DIV13 to DIV16), highlighted in yellow. The dotted line indicates the normalized maximum value of 1.0. g) Normalized P*_young_* TUJ1⁺ and SOX2⁺ subpopulations across development: TUJ1⁺ subpopulation reached peak value at early stages (DIV4-7) when SOX2⁺ expression was lowest, while SOX2⁺ subpopulation increased toward mid-to-late stages. Error bars: SD in b, d, and e; SEM in f and g.

We observed clear age- and stage-dependent differences in marker composition between P*_young_* and P*_old_* cultures (Figure 3.a, 3.b). Across the combined dataset, two-way ANOVA showed significant effects of both *in vitro* developmental stage and age group (both p < 0.0001). The total SOX2⁺ cells were higher in P*_young_* than P*_old_* from DIV7 to DIV14; in P*_young_* it increased over this period, while in P*_old_* it stayed lower and roughly constant.

The two SOX2⁺ cell subpopulations exhibited distinct patterns of behavior. The SOX2⁺/TUJ1⁺ fraction was high at DIV1 in both age groups (no significant P*_young_* - P*_old_* difference), but declined sharply by DIV7. From DIV7 onward, P*_young_* cultures retained a higher SOX2⁺/TUJ1⁺ fraction than P*_old_* at matched stages (DIV7, n(P*_young_*) = 1905 and n(P*_old_*) = 762; DIV10-11, n(P*_young_*) = 1943 and n(P*_old_*) = 2285; and DIV14-16, n(P*_young_*) = 1679 and n(P*_old_*) = 692). The SOX2⁺/vimentin⁺ fraction followed another pattern. Two-way ANOVA revealed significant effects of developmental stage and age group, as well as stage-age group interaction (all p < 0.0001). In P*_young_* it increased steadily from DIV7 to DIV14; in P*_old_* it increased from DIV7 to DIV10 and then dropped by DIV14. Overall, P*_young_* cultures maintained a larger and more dynamic population of immature SOX2⁺ cells, whereas P*_old_* cultures exhibited reduced immature SOX2⁺, consistent with P*_old_* being developmentally more advanced.

We next examined whether quantitative differences in soma size accompanied these differences in cellular composition. The SICM images in Figure 2 indicated an increase in soma size during *in vitro* development, so we quantified soma dimensions and three-dimensional morphology in Figure 3.c, 3.d, and Figure S3. Soma width and length increased proportionally, so cells increased overall rather than elongating along a single axis. At DIV13, P*_young_* somata had a larger area, defined as the two-dimensional soma footprint in the image plane, than P*_old_* somata (p < 0.05). P*_young_* somata also exhibited a larger surface area that captures the three-dimensional contours and protrusions of the soma surface (p < 0.001, Figure S3.a). By DIV21, the area difference was less pronounced, but P*_young_* somata had a significantly larger volume than P*_old_* (p < 0.05; n = 14 cells per group, from two independent coverslips). Age-dependent differences in soma size therefore persisted throughout *in vitro* development, shifting from an area difference at DIV13 to a volume difference at DIV21.

### 2.4 Stage-dependent soma remodeling in P*young* cultures

We quantified soma size in P*_young_* cultures from SICM height maps at DIV1, DIV10, DIV13, DIV15, and DIV21 (Figure 3.e), extracting thickness, area, surface area, volume, and surface roughness. One-way ANOVA showed a significant effect of developmental stage on every parameter (all p < 0.001). Area, surface area, thickness volume, and roughness increased progressively from the earliest stages to DIV15, then decreased by DIV21. The soma volume increased from 277 ± 82 µm³ at DIV1 to 821 ± 209 µm³ at DIV15, then decreased to 565 ± 289 µm³ at DIV21. Surface roughness increased from 1.10 ± 0.20 µm at DIV1 to 1.19 ± 0.22 µm at DIV15, then declined to 0.71 ± 0.38 µm at DIV21 (n = 14).

P*_old_* cultures, by contrast, changed little once established. Between DIV13 and DIV21, two-way ANOVA revealed no significant change in thickness, area, surface area, volume, or roughness (all p > 0.05). P*_young_* somata increased in size and then partly shrank during *in vitro* development, while P*_old_* somata remained comparatively stable.

### 2.5 Morphometry and cellular composition co-vary

We next compared SICM-derived morphometric parameters with fluorescence-based cellular composition across P*_young_ in vitro* development (Figure 3.f, 3.g). We normalized soma thickness, area, volume, roughness, and marker-positive subpopulations to compare their relative temporal trajectories across *in vitro* development. Because we obtained fluorescence and SICM measurements from separate cells, this comparison reflects population-level temporal alignment rather than cell-by-cell correlation.

Area, volume, and thickness increased from early stages, peaking around DIV13-DIV16, coinciding with the highest abundance of SOX2⁺ cells. Surface roughness is less uniform, with relatively high values early on and lower values later. These trends indicate that soma enlargement and SOX2-associated cellular composition were temporally aligned during development.

We also compared normalized TUJ1⁺ and SOX2⁺ subpopulations in P*_young_* cultures (Figure 3.g). TUJ1⁺subpopulations were highest at early stages, while SOX2⁺ subpopulations increased toward mid-to-late stages. TUJ1⁺ cells were not quantified at DIV21 because the cultures were too confluent for reliable counting. Taken together, soma size measured from SICM and cellular composition obtained from fluorescence microscopy varied across the same stages of P*_young_ in vitro* development.

### 2.6 Composition and structure of glial cells

We next examined glial cell morphology and cellular composition in P*_young_* and P*_old_* cultures. Correlative brightfield and SICM imaging revealed distinct three-dimensional morphologies of neurons and glial cells. Neurons displayed compact cell bodies with steep height gradients, whereas glial cells exhibited broader and flatter morphologies with extended lamellar regions, as confirmed by the corresponding height profiles (Figure 4.a).

**Figure 4.**
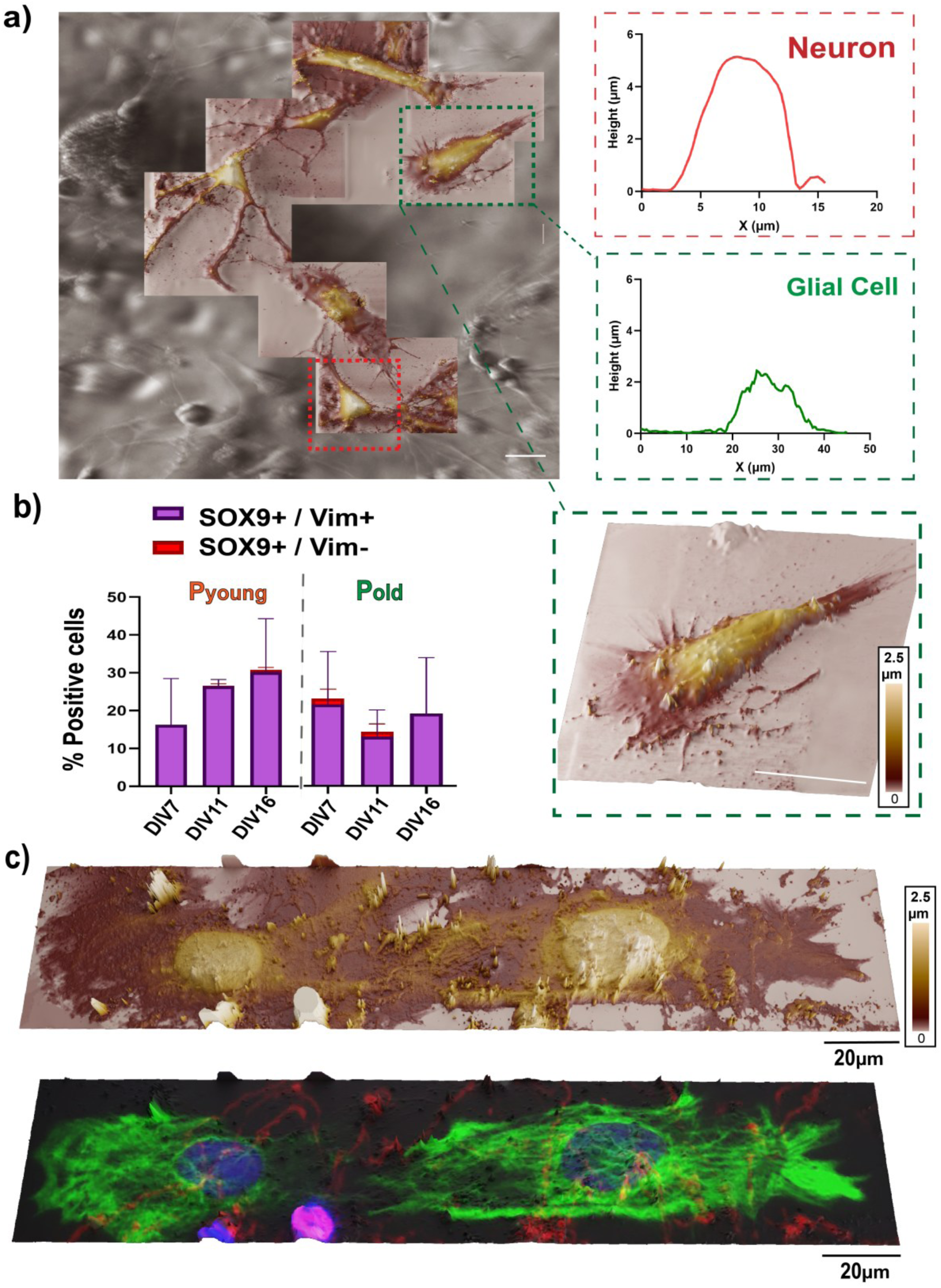
Glial cell composition and label-free structural discrimination. a) Correlative brightfield and SICM imaging of neurons and glial cells with corresponding height profiles. Neurons displayed compact somata and steep height gradients, whereas glial cells had broader, flatter morphology; scale bar = 20 µm. b) Proportions of SOX9⁺/Vim⁻ and SOX9⁺/Vim⁺ cell subpopulations in P*_young_* and P*_old_* cultures during *in vitro* development. The SOX9⁺/Vim⁻ astroglia subpopulation increased over time in P*_young_* cultures, whereas in Pold cultures it peaked at DIV7 and then declined, with lower values at DIV11 and DIV16. SOX9⁺/Vim⁺ (immature/reactive astrocytes) accounted for a small proportion of the cell culture, with the most prominent values in P*_old_* cultures at DIV7. c) Correlative SICM (upper panel) and fluorescence imaging (lower panel) of the same region representing glial cells, combining marker-based cellular identification with label-free three-dimensional surface morphology. Vimentin (green), TUJ1 (red), DAPI (blue), Scale bar = 20 µm

To assess glial cell composition, we immunostained for SOX9, a marker of astroglial identity, and vimentin, a marker of immature or reactive glia. SOX9**⁺** cells were divided into vimentin-negative (SOX9⁺/Vim⁻) and vimentin-positive (SOX9⁺/Vim⁺) subpopulations (Figure 4.b, Figure S4). At early stages (i.e., DIV7), P*_old_* cultures contained a greater proportion of SOX9⁺ astroglial cells than P*_young_*, with most of these cells displaying a SOX9⁺/Vim⁺ phenotype. As cultures aged *in vitro* (i.e., DIV 11-16), the SOX9⁺/Vim⁻ astroglial subpopulation expanded in P*_young_* cultures, whereas in P*_old_* cultures, the SOX9⁺/Vim⁻ subpopulation peaked at DIV7, then decreased at DIV11, followed by partial recovery by DIV16. At DIV16, P*_young_* cultures displayed a larger proportion of SOX9⁺ astroglia than P*_old_* cultures. The SOX9⁺/Vim⁺ population remained as a minor proportion in both age groups and was most prominent in P*_old_* cultures at DIV7 (Figure 4.b).

Correlative imaging of the same regions using two modalities, SICM and fluorescence microscopy, provided complementary information on cellular identity and structure (Figure 4.c). Fluorescence imaging identified distinct cell subpopulations based on marker expression, whereas SICM resolved the three-dimensional surface architecture of the corresponding cells, including broad lamellar regions and irregular surface topography. These observations demonstrate that label-free SICM of cellular morphology can complement fluorescence-based identification.

### 2.7 Injury response differs in P*young* and P*old* cultures

We compared injury responses in cell cultures following a mechanical transection (a scratch lesion; see Methods). We made lesions at DIV7, DIV10, and DIV14, then fixed cultures 24 or 48 h later for fluorescence and SICM imaging (Figure 5; Figures S5a, S5b and S6). Fluorescence imaging showed TUJ1⁺ neuronal processes and vimentin⁺ glia within and surrounding the lesion (Figure 5.a). Elongated vimentin⁺ cells lined up and oriented along the lesion in both groups, but their share of lesion coverage differed by age.

**Figure 5.**
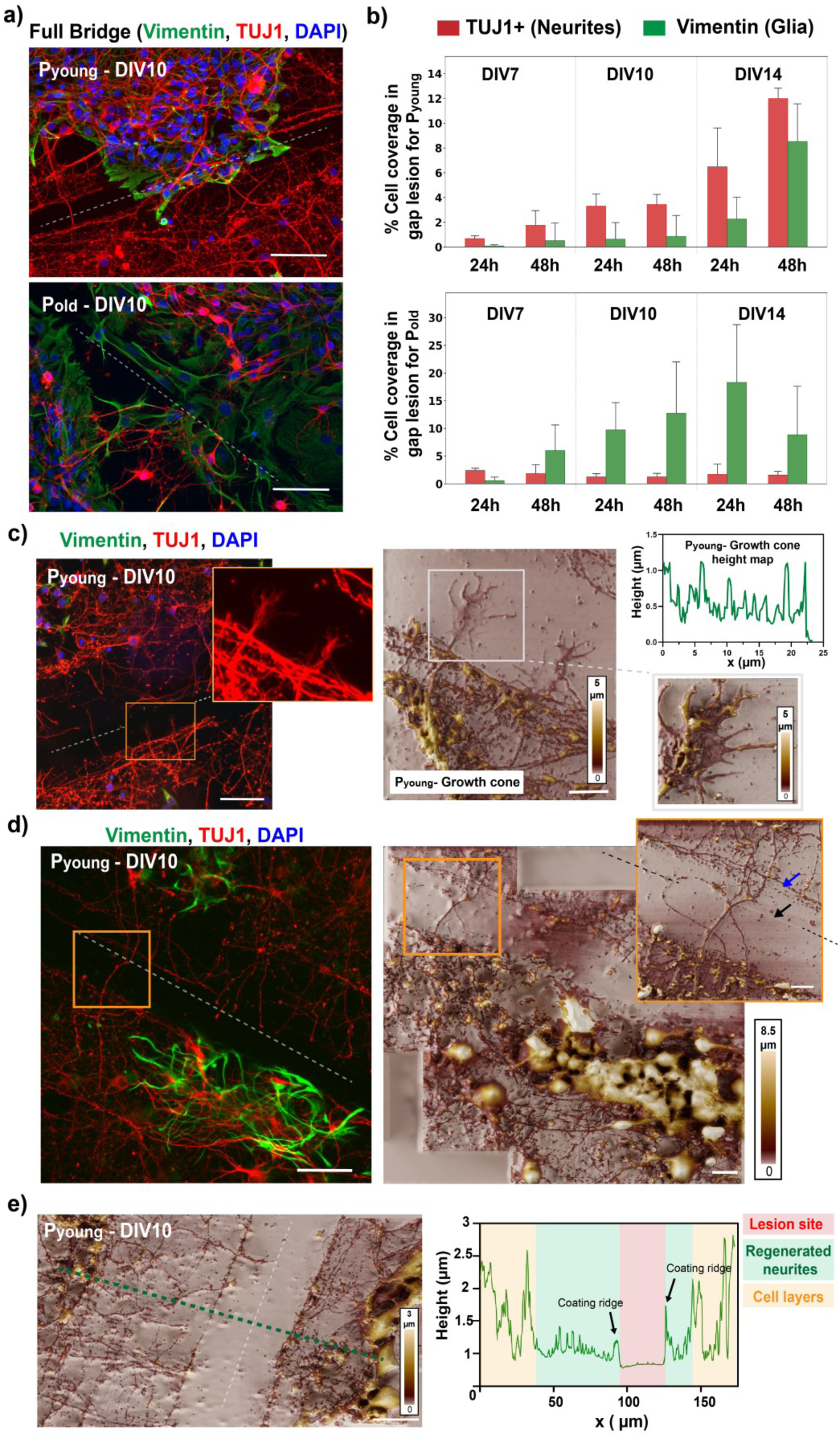
Fluorescence microscopy and label-free SICM imaging reveal age-dependent remodeling during lesion repair. Fluorescence microscopy: vimentin (green) labeled reactive or elongated glial cells, TUJ1 (red) labeled neuronal processes, and DAPI (blue) marked nuclei. a) P*_young_* and P*_old_* cultures at DIV10, 48 h-post injuries, respectively: P*_young_* cultures displayed neurite-rich bridges across the lesion, whereas P*_old_* cultures showed predominant accumulation of vimentin⁺ glial cells. Scale bar, 50 µm. b) TUJ1⁺ neurite and vimentin⁺ glial cell coverage of the lesion gap in P*_young_* (upper panel) and P*_old_* (lower panel) at DIV7, DIV10, and DIV14, at 24 h- and 48 h post-injury (one-way ANOVA, p < 0.001). c) Correlative fluorescence microscopy and SICM of the boxed lesion-border region in P*_young_* cultures at DIV10; SICM height maps showed growth cone shape and height. Scale bar, 10 µm. d) Correlative fluorescence microscopy and SICM imaging of the lesion site at DIV10; the magnified region showed TUJ1⁺ neuronal processes aligned parallel to the scratch-induced coating ridges (blue arrow) and the transection groove (black arrow). Scale bar, 10 µm. e) SICM image of the lesion with a cross-sectional height profile showing the coating ridges and three distinct regions: central lesion site, regrown neurites, and surrounding cell layers. Scale bar, 20 µm.

In P*_young_* cultures, TUJ1⁺ neurite coverage of the lesion gap increased with days *in vitro*, beginning with 0.68 ± 0.23% at DIV7 (24 h post-lesion) to 12.00 ± 0.83% at DIV14 (48 h post-lesion) (Figure 5.b), while vimentin⁺ glial coverage remained relatively low. In P*_old_* cultures, we observed the opposite pattern. The lesion was predominantly occupied by vimentin⁺ glia, reaching 9.75 ± 4.90% at DIV10 (24 h-post lesion) and 18.29 ± 10.44% at DIV14 (24 h-post lesion), while TUJ1⁺ neurite coverage of the lesion remained limited across all time points. In summary, P*_young_* cultures exhibited predominantly neurite-based lesion repair, while P*_old_* cultures showed mainly glial accumulation at the lesion site.

Correlative fluorescence microscopy and SICM imaging compiled structural detail at the lesion border. In P*_young_* cultures, SICM resolved growth cone and filopodia-like surface structures near the lesion site, along with their height profiles (Figure 5.c). We observed similar but smaller, simpler growth cone-like structures in P*_old_* cultures (Figure S5.b). Because these came from representative images rather than systematic counting and analysis, we treated them as qualitative features of the lesion response.

SICM also resolved the physical topography of the lesion site. Height maps showed the transection groove and parallel ridge-like structures formed when scratching partially removed and displaced the coating. TUJ1⁺ processes were observed along these ridges (Figure 5.d, 5.e). Three-dimensional height maps distinguished the cell bodies, the layer of regrown cellular extensions, and the remaining lesion gap. SICM therefore captured both cellular structures at the lesion border and the substrate topography produced by mechanical transection.

## 3. Discussion

A central challenge in neuroregeneration research is linking molecular cell identity to the physical state of cells and tissues. This study addressed this gap by showing that measurable changes in three-dimensional morphology and injury response accompany developmental changes in cellular composition. SICM can capture these nanoscale structural features label-free, providing information that molecular markers alone cannot. Regeneration should therefore be considered not only a molecular process but also a physical reorganization of cells, cellular processes, and surrounding tissue architecture.

Because regeneration reactivates developmental processes that decline as the CNS matures^2,3,5^, we tracked markers associated with developmental, neuronal, and glial states. Fluorescence analysis showed that P*_young_* cultures retained a larger and more dynamic pool of immature SOX2⁺ cells, whereas P*_old_* cultures contained a greater proportion of SOX9⁺/vimentin⁺ glial cells. These findings align with previous reports of stage-dependent SOX2 and SOX9 expression in the postnatal opossum cortex51 and suggest that Pyoung cultures retain a more immature state, while Pold cultures shift toward glial cells. This pattern is consistent with the age-dependent transition toward gliogenesis^41,42^, which in the opossum begins around P16-P18^55^.

A key finding of the multimodal analysis was the temporal alignment between SICM-derived morphometry and fluorescence-based cellular composition. P*_young_* and P*_old_* cultures differed not only in cellular composition but also in soma morphology. In P*_young_* cultures, soma size increased and subsequently declined during *in vitro* development, while P*_old_* soma remained relatively stable between DIV13 and DIV21. Notably, in P*_young_* cultures, soma size peaked around DIV13-DIV16, coinciding with a high proportion of SOX2⁺ cells. Thus, the period associated with the greatest change in molecular composition also showed pronounced changes in cell morphology. The increase and subsequent decline in P*_young_* soma size may reflect a transition from active cellular growth to morphological stabilization, consistent with previous reports that synaptic density in cortical cultures peaks around DIV14 and then declines ^56^. Surface roughness followed a similar course. Because actin-dependent cytoskeletal remodeling^57^ can influence membrane surface roughness, the observed changes may reflect a transient increase in membrane dynamics. However, SICM-measured surface roughness does not directly measure actin. These temporal associations do not prove causation, but they demonstrate that changes in cellular composition and morphology occur over similar developmental stages.

The lesion response showed the same split. Across developmental time *in vitro*, P*_young_* cultures showed greater coverage of the lesion with TUJ1⁺ neurites, while P*_old_* cultures exhibited limited neurite coverage and a predominance of vimentin⁺ glia. The glia-dominated response in P*_old_* cultures is consistent with previous studies in which reactive glia contribute to scar formation and limit tissue repair^4,11,15^. At the lesion border, SICM resolved growth cones and filopodia-like structures, as well as their height profiles. Given the central role of growth cones and cytoskeletal remodeling in axonal growth^9,10,13,14^, this highlights the potential of SICM to study lesion-associated surface morphology. However, because these features were not systematically quantified, they should be considered a qualitative observation.

SICM also showed, at higher resolution, that the mechanical lesion changed the substrate itself, leaving coating ridges parallel to the scratch, with TUJ1⁺ processes aligned along them. Alignment in this assay therefore reflects both cellular repair and the physical shape of the scratched surface, and possibly chemical guidance cues such as netrins^58^, semaphorins^59^, and ephrins^60^ that we did not measure. This is a limitation of *in vitro* scratch injuries and argues for more controlled lesioning, for example axotomy with micro-/nanopositioning, laser^61^ or microfluidics^62^.

Together, these results extend earlier transcriptomic^33,34^ and proteomic^32,63^ studies of stage-dependent molecular differences between regeneration-permissive and non-permissive opossum CNS by showing that those differences come with measurable differences in cell shape. SICM adds a label-free structural layer to molecular and marker-based regeneration analyses.

Several limitations should be considered. First, the relationship between cellular composition, SICM-derived morphometry, and lesion repair is correlative. Targeted functional studies are therefore needed to determine whether glial composition, SOX2⁺ cell populations, cytoskeletal remodeling, or substrate topography directly influence repair. Second, dissociated cortical cultures do not reproduce the full tissue architecture, extracellular matrix, immune environment, vascular context, or long-range connectivity of the intact CNS. Third, we performed SICM morphometry on fewer biological preparations than fluorescence quantification, limiting population-level inference. Finally, SICM measurements were performed on fixed samples, enabling correlative imaging with immunofluorescence at selected cellular locations while avoiding the technical challenges of live fluorescent labeling and phototoxicity. However, this approach limited the assessment of dynamic morphological changes. Future studies could combine live-cell SICM with subsequent fixation and immunostaining, linking temporal changes in cell structure to molecular identity.

Future work should extend this to live-cell SICM and mechanical mapping. SICM uses non-contact live imaging and can estimate local mechanical properties^29,64^. Because mechanical cues influence nervous system development, neuronal growth, and glial pathology^65,66^, live structural and mechanical measurements could show whether the shape differences we see merely accompany regeneration-linked responses or actively help drive them.

## 4. Conclusion

In postnatal opossum cortical cultures, developmental stage was associated with distinct changes in marker-defined cellular composition, three-dimensional soma morphology, and injury response. Pyoung cultures showed greater morphological remodeling and predominantly neurite-based lesion repair, whereas Pold cultures exhibited more stable soma morphology and a stronger glial response. The temporal association between cellular composition and SICM-derived morphometry demonstrates how label-free three-dimensional imaging can complement fluorescence-based characterization of developmental state. Together, these findings highlight SICM as a valuable quantitative complement to fluorescence microscopy for linking developmental cell-state changes with three-dimensional structural differences across changes in CNS regenerative capacity.

## 5. Materials and Methods

### 5.1 Animal model and experimental cohorts

Experiments used the South American gray short-tailed opossum (Monodelphis domestica) as a developmental model of CNS regeneration. The Local Veterinary Service, the Ethical Committee of the Department of Biotechnology, University of Rijeka (approval dated January 4 2017), and the National Ministry of Health (Permit No. 1FF80.N.9Q3) approved all procedures, which were conducted in accordance with European Directive 2010/63/EU. In 2023, the Department of Biotechnology was renamed the Faculty of Biotechnology and Drug Development, University of Rijeka.

Animals of both sexes were used. We defined two cohorts to study age-dependent differences in regenerative potential, including young pups (P3-P6), the regeneration-permissive stage, and older pups (P16-P18), the non-permissive stage. We maintained animals at the University of Trieste animal facility under controlled conditions (27-28 °C, 50-60% humidity) with a 12 h light/dark cycle and *ad libitum* access to food and water.

### 5.2 Primary Dissociated Cortical Cultures

Primary cortical neuronal cultures were isolated from P3-P6 and P16-P18 opossum pups based on our previously established and described protocol^45^. Briefly, glass coverslips (12 mm) were acid-washed, sterilized, and coated with poly-L-ornithine (50 µg/mL, Sigma-Aldrich) followed by laminin (2 µg/mL, Sigma-Aldrich) prior to cell plating.

Following euthanasia, brains were rapidly extracted and placed in ice-cold oxygenated dissection solution. Cerebral cortices were isolated bilaterally, and meninges, olfactory bulbs, and subcortical structures were removed. Tissue was enzymatically dissociated using age-adapted trypsin digestion (0.5% w/v trypsin in phosphate buffer solution, PBS for P3-P6; 2.5% w/v trypsin in PBS for P16-P18, Santa Cruz Biotechnology) at 32.5°C, followed by mechanical trituration in a solution containing DNAse I (10 µg/ml, Sigma-Aldrich), trypsin inhibitor (1 mg/ml, Santa Cruz Biotechnology) and bovine serum albumin (BSA, 1% w/v, Pan-Biotech) in Hank’s Balanced Salt Solution w/o Ca^2+^ and Mg^2+^ (HBSS, Pan-Biotech). Cell suspensions were purified by 5% BSA in HBSS gradient centrifugation and resuspended in plating medium containing DMEM supplemented with 10% fetal bovine serum (FBS) and 1% penicillin/streptomycin (all from Pan-Biotech). Cells were briefly pre-plated to reduce fibroblast contamination, counted with a hemocytometer, and seeded at a density of 5 × 10⁴ cells per well. For each isolation, cortical tissue from three to four pups, depending on pup size, was pooled before dissociation. The resulting cell suspension was distributed across multiple coverslips, which were maintained for analysis at different days in vitro (DIV).

After 24 hours, we partially replaced the medium with serum-free neuronal medium consisting of Neurobasal medium supplemented with B27 (both from Thermo Fisher Scientific), L-glutamine, and 1% penicillin/streptomycin (both from Pan-Biotech). Cultures were maintained at 32°C in a humidified incubator with 5% CO₂ and 95% relative humidity. Half of the medium was replaced weekly. In accordance with the 3R principles^43,44^, the number of animals was minimized by pooling cortices from all pups per culture preparation.

Early postnatal (P3-P6) cultures remained viable for more than one month *in vitro* and exhibited high neuronal purity (>93%). Late postnatal (P16-P18) cultures showed reduced neuronal proportion (∼80%) and increased glial representation (∼20%), consistent with developmental progression toward gliogenesis^45^. Cells were analyzed at defined *in vitro* stages representing early (∼DIV7), intermediate (DIV9-11), mid-to-late (DIV13-16), and late development (DIV17-30). A wound healing (scratch) assay was performed according to previous studies^67^. Mechanical lesions were made by drawing a fine-tipped tweezer (Dumont) across the center of the coverslip, severing the differentiated processes and producing a scratch approximately 100 µm wide. Cultures were lesioned at DIV7, DIV10, or DIV14 and fixed 24 or 48 h after injury for fluorescence and SICM analysis.

In the original culture series, coverslips examined at different post-injury timepoints originated from the same pooled cell preparation. Non-injured cultures were maintained separately and originated from different pooled preparations.

### 5.3 Immunofluorescence Analysis

To assess lineage composition and cellular identity, cultures were fixed at defined *in vitro* stages and processed for immunocytochemistry. Cells were fixed for 20 min with 4% w/v paraformaldehyde supplemented with 200 mM sucrose, pH 6.9 (both from Sigma-Aldrich), saturated with glycine (0.1 M, Sigma-Aldrich), permeabilized with Triton X-100 (0.1% v/v, Sigma-Aldrich) in PBS, and blocked with 0.5% w/v BSA (Pan-Biotech) in PBS for 30 min to prevent nonspecific antibody binding. Primary antibodies targeting neuronal and glial markers were applied, including βIII-tubulin (TUJ1), SOX2, SOX9, vimentin, and GFAP. Primary antibodies were incubated for 1 h at room temperature, followed by washing and applying appropriate fluorescently labeled secondary antibodies for 30 min. A full list of antibodies is provided in Tables S1 and S2. Nuclei were counterstained with 4’,6-diamidino-2-phenylindole (DAPI, 300 nM, Thermo Fisher Scientific in PBS). Fluorescence imaging was performed using an inverted fluorescence microscope (Olympus IX83, USA) equipped with differential interference contrast and fluorescence optics. Images were acquired using 10× (0.4 NA) and 20× (0.5 NA) objectives. For each sample, a minimum of 10 z-stack frames were collected with objective-specific slice spacing, and maximum intensity projections were generated.

Image processing and quantitative analysis were performed using dedicated microscopy software (CellSens, USA) and ImageJ/Fiji software. Lineage proportions were calculated as the percentage of marker-positive cells relative to total DAPI⁺ nuclei. Co-expression analysis was performed to determine proportions of SOX2⁺ and TUJ1**⁺** and similarly for SOX9⁺ and vimentin**⁺** cell populations.

For lesion-coverage analysis, the cut area was manually isolated in Fiji/ImageJ. TUJ1⁺ neurite and vimentin⁺ glial fluorescence within this region was quantified, and coverage was calculated as the fluorescent-marker-positive area divided by the total lesion area × 100.

For fluorescence quantification, multiple coverslips were examined at the investigated developmental stages and age groups. Several microscopy fields were acquired from each coverslip, and marker-positive cells were counted relative to the total number of DAPI-positive nuclei. Cell counts from the examined coverslips and microscopy fields were pooled within each experimental condition to determine the proportions of marker-positive cells. The total number of cells analyzed (n) is reported for each condition.

### 5.4 Scanning ion conductance microscopy (SICM)

Topographic imaging used a non-contact SICM system based on a previously described design^27^. The system comprises a controller for feedback control of fine movement, a custom-built Z-actuator, a stepper motor for coarse movement, and an XY scanner (P-517 CL Piezo NanoPositioner, Physik Instrumente, 100 µm travel range), mounted on an Olympus IX71 inverted microscope. Glass nanopipettes were fabricated to tip diameters of 120-150 nm, giving an estimated lateral resolution of about 1.5 × pipette diameter and an axial resolution below 5 nm^28^. Borosilicate glass (Sutter Instruments, USA) was pulled with a CO₂ laser puller (Model P-2000, Sutter Instruments, USA).

Imaging used hopping mode to avoid contact with the cell surface. A bias voltage of −200 mV was applied between the pipette and bath electrodes. Ion current was sampled at 1 MHz, with baseline currents typically 4-5 nA. Surface detection was set to a 1% drop in ion current (setpoint 99% of baseline). The vertical approach rate was 30 µm/s, and the probe retracted 7 µm between hops, at an effective hopping rate of ∼75 Hz. Scans covered 60-80 µm areas at 256 × 256 or 128 × 128 pixels.

SICM height maps were analyzed in Gwyddion after background leveling. Individual somata were manually segmented, excluding incomplete or overlapping cells. Thickness, projected area, 3D surface area, volume, and Sa roughness were calculated within the selected soma region. The same analysis procedure was applied to all images, with no smoothing or filtering before quantification. Larger fields of view were stitched from adjacent scans in ImageJ/Fiji.

SICM morphometry was performed on cells from two separate coverslips for each developmental stage and age group. Morphological parameters were quantified for individual somata, and the measurements were combined to characterize cell morphology within each condition. The reported cell number (n) represents the total number of somata analyzed.

### 5.5 Statistical analysis

Fluorescence-based cellular composition and SICM morphometry were measured in separate cells at corresponding developmental stages. For fluorescence analysis, cell counts from multiple coverslips belonging to the same age group, developmental stage, and experimental condition were pooled, whereas SICM morphological parameters were quantified from individual somata. Data are presented as mean ± standard deviation unless otherwise stated, and sample sizes are reported in the corresponding sections. For comparisons among three or more groups, we used one-way ANOVA followed by Tukey post-hoc testing for normally distributed data with equal variances. We performed two-group comparisons using unpaired t-tests. We used two-way ANOVA followed by Tukey post-hoc testing to assess the effects of developmental stage, age group, and their interaction. We set statistical significance at p < 0.05. We performed analyses using GraphPad Prism 10.3.

## Author contributions

Z. A.: conceptualization, investigation, formal analysis, visualization, writing original draft, writing review and editing.

M. P.: investigation, formal analysis, writing review and editing.

M. M.: conceptualization, resources, writing review and editing.

J. B.: conceptualization, resources, supervision, writing review and editing.

G. E. F.: conceptualization, resources, supervision, writing review and editing.

## Data Availability

The data supporting this article are provided in the Supplementary Data. Additional data are available from the corresponding authors upon reasonable request.

## Ethics Approval

All procedures involving South American gray short-tailed opossums (*Monodelphis domestica*) were approved by the Local Veterinary Service, the Ethical Committee of the Department of Biotechnology, University of Rijeka (approval dated January 4, 2017), and the National Ministry of Health (Permit No. 1FF80.N.9Q3), in accordance with European Directive 2010/63/EU. The experiments were additionally approved by the EPFL Animal Research Ethics Committee (AREC, approval no. AREC000016). In 2023, the Department of Biotechnology was renamed the Faculty of Biotechnology and Drug Development, University of Rijeka. All neuronal samples were received at EPFL as fixed specimens, and no experiments involving live animals were conducted at EPFL.

## Acknowledgments

This project received funding from the European Union’s Horizon 2020 research and innovation program under the Marie Skłodowska-Curie grant agreement No 945363. G.E.F. and Z.A. also acknowledge the Swiss Commission for Technology and Innovation under grant CTI-18330.1, and the EPFL Center for Imaging under grant No. 563292. We also acknowledge the Croatian Science Foundation (HRZZ/CSF) grant IP-2016-06-7060 to M.M. and support from the University of Rijeka (uniri-prirod-18-290-1463 to J.B.; uniri-biomed-18-258-6427 to M.M.).

