## supplementary information for "Multimodal imaging links molecular identity with nanoscale-resolved 3D cell morphology in opossum cortical cultures"

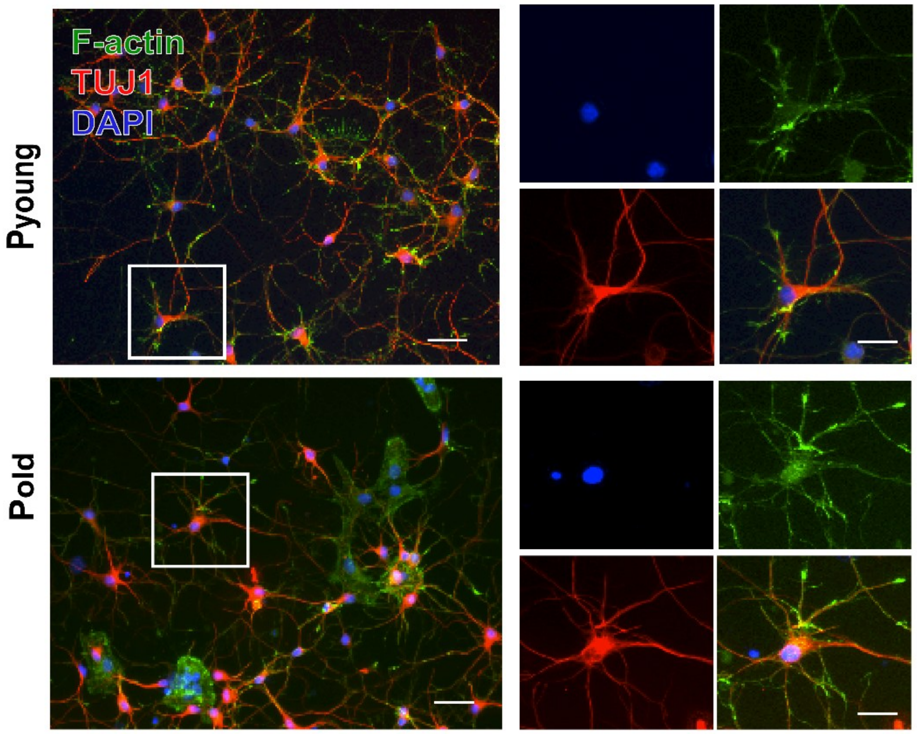


**Figure S1.** At DIV7, both $P_{young}$ and $P_{old}$ neurons exhibit extensive neurite outgrowth and connections, including fine processes such as spines, as visualized by microtubules (TUJ1, red) and microfilaments (F-actin, green). F-actin was labelled with phalloidin i-Fluor 488 (Abcam, Cat# ab176753), diluted 1:100 and incubated 30 min at room temperature together with the secondary antibodies. Cell nuclei are counterstained with DAPI (blue). Scale bar, 50 µm.

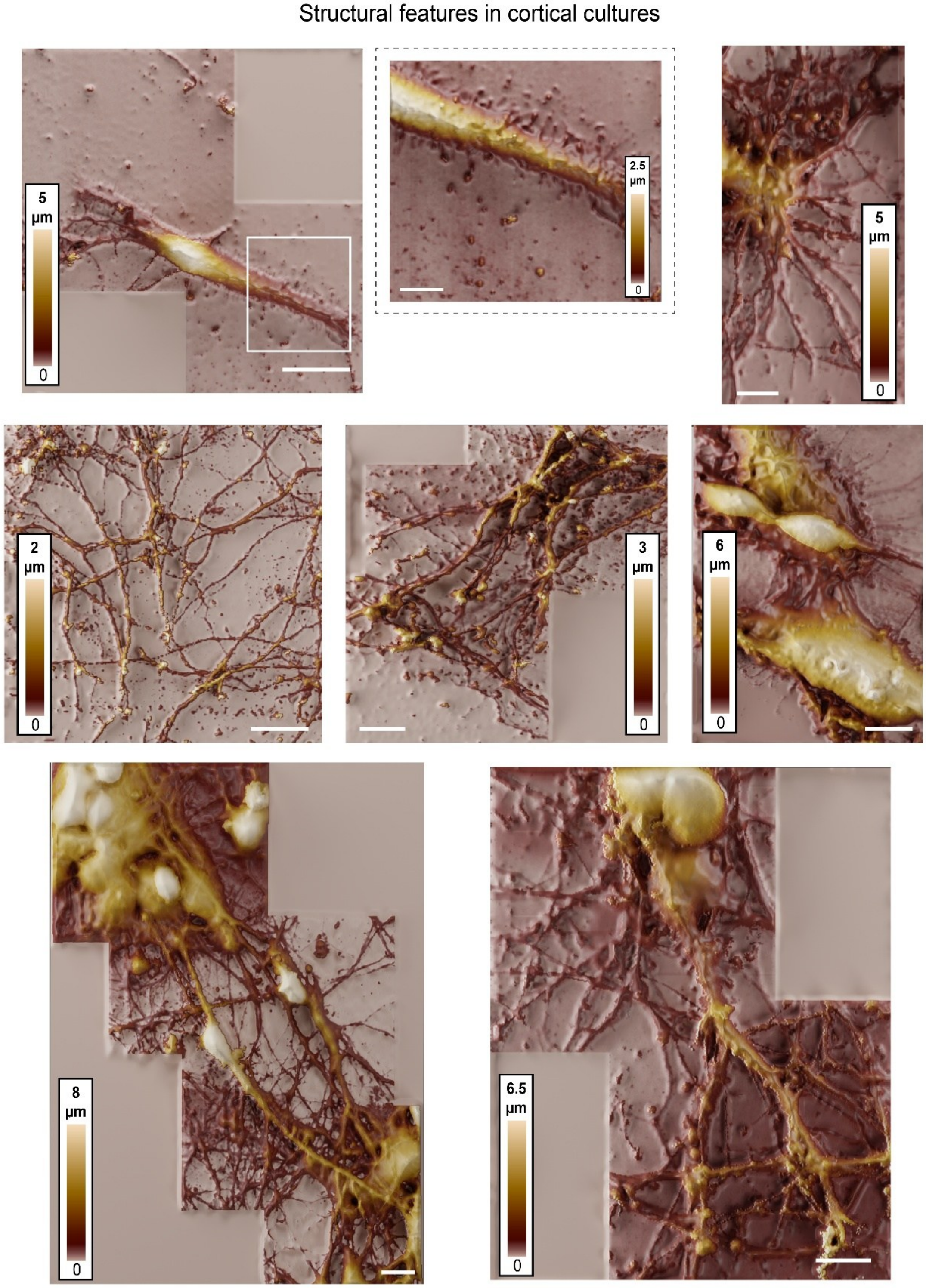


**Figure S2.** Representative SICM images showing diverse structural features in cortical cultures, including soma, neuronal networks, glial cells, and fine processes such as neurites and filopodia. Scale bar, 10 µm.

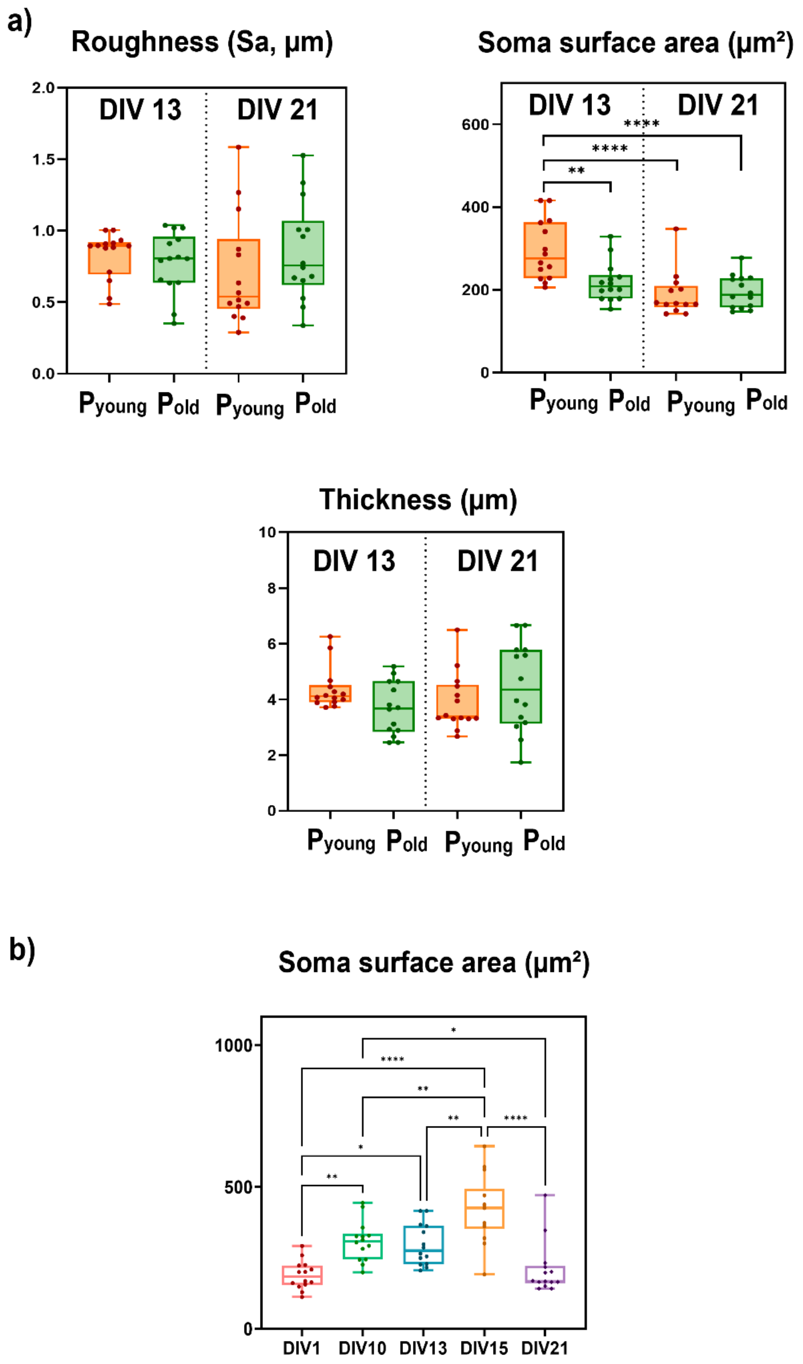


**Figure S3.** Statistical morphometric analysis of SICM images; a) Analysis of roughness, thickness and surface area between $P_{young}$ and $P_{old}$ cultures. b) Analysis of soma surface area in $P_{young}$ cultures (one-way ANOVA, $p < 0.001$; n: DIV1 = 14, Tukey post-hoc; *$p < 0.05$, **$p < 0.01$, ***$p < 0.001$, ****$p < 0.0001$).

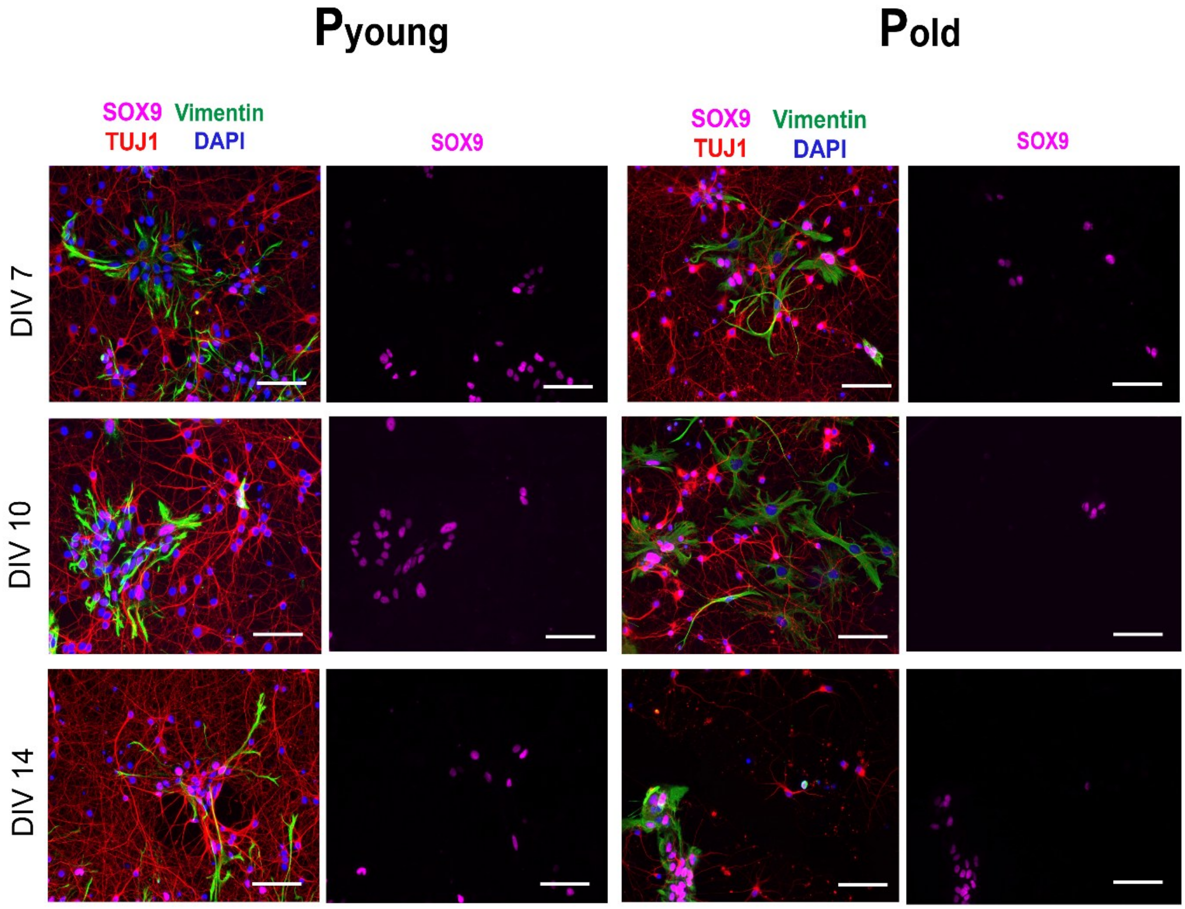


**Figure S4.** Fluorescence microscopy images of SOX9$^+$ subpopulations in $P_{young}$ and $P_{old}$ cultures across *in vitro* development. SOX9 (pink) marks nuclei of glial progenitors and astrocytes, vimentin (green) labels cytoskeletal filaments in populations including radial glia and reactive astrocytes, and TUJ1 (red) labels neuronal cell bodies and neurites. Scale bar, 50 µm.

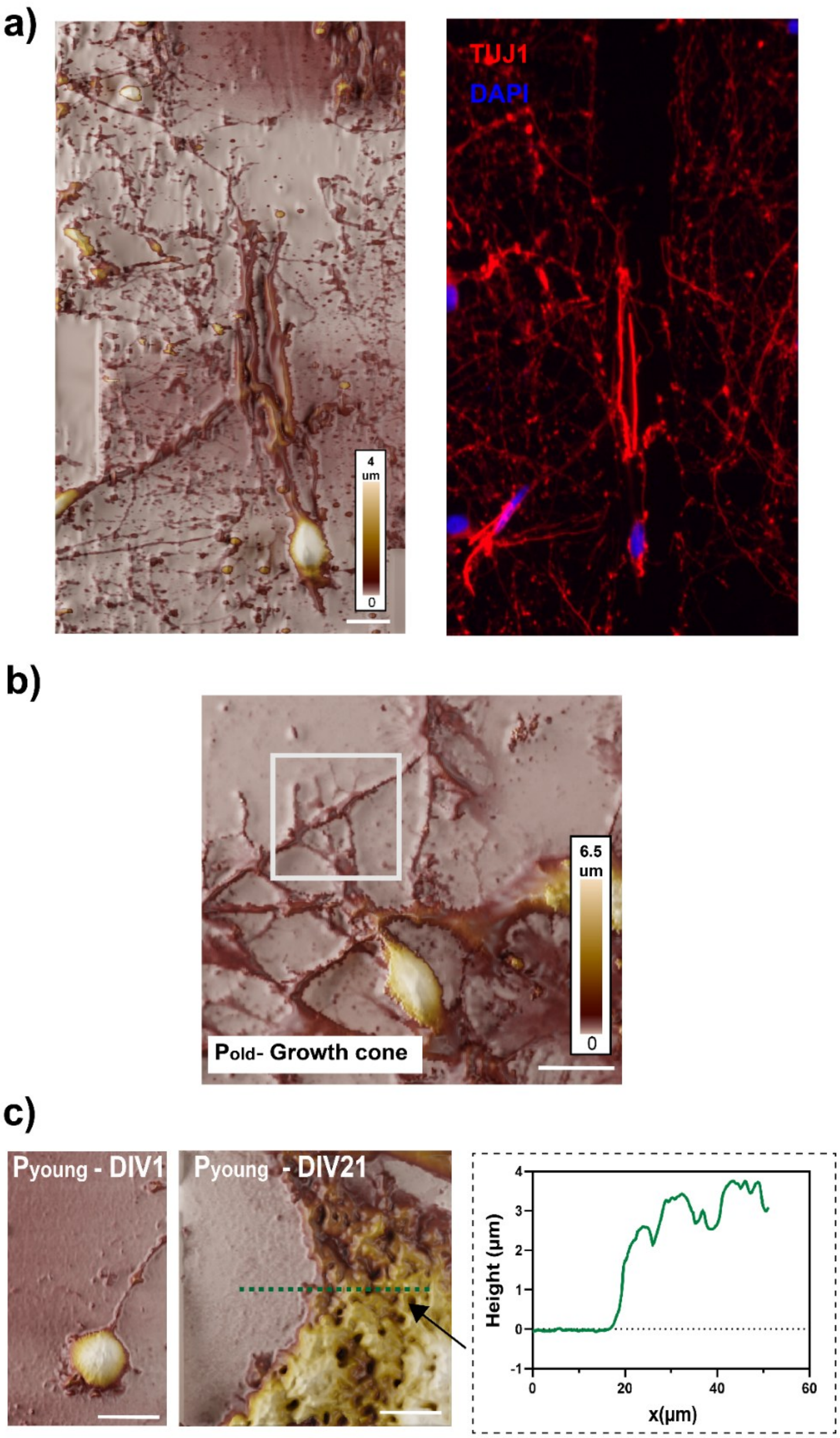


**Figure S5.** Analysis of the lesion site. a) Correlative SICM and fluorescence microscopy images of the same region at the lesion border, showing how the two modalities provide complementary structural and molecular information. SICM resolves three-dimensional surface architecture (scale bar, 10 µm), including membrane topology and fine cellular extensions, while fluorescence microscopy (scale bar, 50 µm) identifies lineage-specific markers, demonstrating the value of combining both approaches for studying regeneration. b) SICM height map image of a $P_{old}$ culture growth cone at DIV10, showing comparatively smaller dimensions and simpler morphology relative to $P_{young}$ cultures; Scale bar = 10 µm. c) After 21 days in culture, cells extended neurites and formed a network over a basal cellular layer, with distinct height differences between the cell bodies, the underlying layer, and the neurite network. Scale bar = 10 µm, x and y in µm.

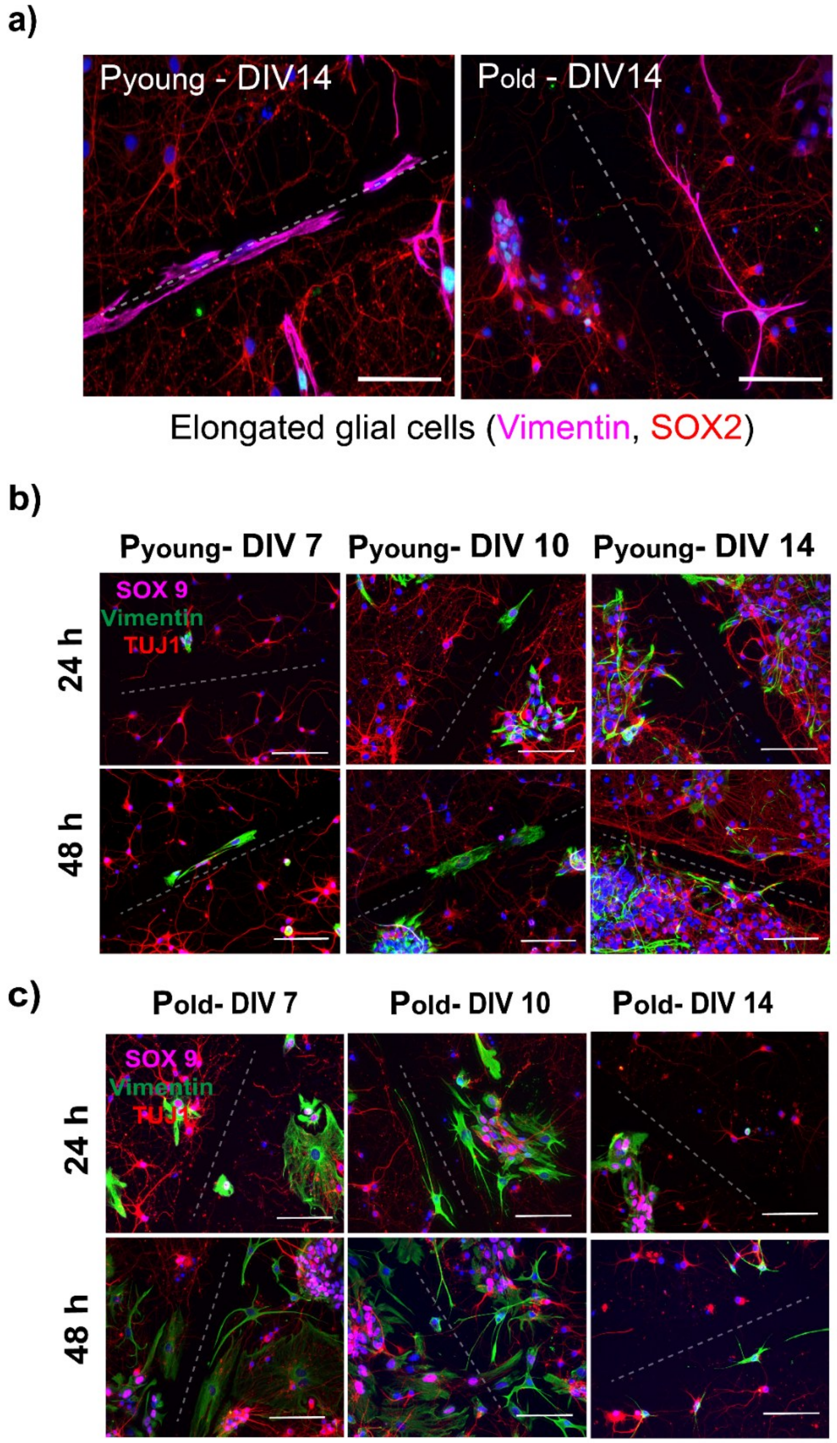


**Figure S6.** Representative fluorescence microscopy images of glial cell involvement during lesion remodeling at different timepoints *in vitro*. a) Elongated glial cells aligned along the lesion axis in $P_{young}$ and $P_{old}$ cultures at DIV14. Scale bar, 50 µm. Vimentin-positive cells (magenta) highlight the elongated glial morphology within the injury region in $P_{young}$ cultures. b)

Fluorescence microscopy images of $P_{young}$ cultures at DIV7, DIV10, and DIV14, 24 h and 48 h post-injury showing cellular composition at the lesion site. SOX9-positive cells (magenta) indicate glial lineage, vimentin (green) marks reactive or elongated glial structures, and TUJ1 (red) labels neuronal processes. Scale bar, 50 µm. c) Corresponding fluorescence microscopy images for $P_{old}$ cultures at DIV7, DIV10, and DIV14, 24 h and 48 h post-injury, showing increased glial accumulation and organization within the lesion region compared to $P_{young}$. Scale bar, 50 µm.

**Table S1 Primary antibodies used in this work.**

| Primary antibody | Host and type | Dilution | Immunogen | Producer Cat # RRID |
|---|---|---|---|---|
| **Anti - SOX2 antibody [9-9-3]** | Mouse monoclonal IgG1 | 1:200 | Synthetic peptide conjugated to KLH derived from within residues 300 to the C-terminus of Human SOX2. | Abcam, Cat# ab79351, RRID: AB_10710406 |
| **Anti - SOX9 antibody [EPR14335-78]** | Rabbit monoclonal IgG | 1:300 | Recombinant fragment within Human SOX9 aa 150-300 (internal sequence). | Abcam, Cat# ab185966, RRID: AB_2728660 |
| **Anti-Tubulin β 3 (TUBB3) Antibody (TUJ1)** | Mouse monoclonal IgG2a | 1:200 | Microtubules derived from rat brain. | Biolegend, Cat# 801201, RRID: AB_10063408 |

| | | | | |
|---|---|---|---|---|
| **Anti - Vimentin antibody [EPR3776]** | Rabbit monoclonal IgG | 1:400 | Synthetic peptide within Human Vimentin aa 400 to the C-terminus (C terminal). The exact sequence is proprietary. | Abcam, Cat# ab92547, RRID: AB_10562134 |
| **Anti - Vimentin antibody** | Mouse monoclonal IgG1 | 1:400 | Aa 350-500. Purified vimentin from pig eye lens. | Abcam, Cat# ab8069, RRID: AB_306239 |
| **Anti – GFAP antibody** | Mouse monoclonal IgG1 | 1:300 | GFAP from pig spinal cord. | Sigma-Aldrich, Cat# G3893, RRID: AB_477010 |

**Table S2 Secondary antibodies used in this work.**

| Secondary antibody | Host and type | Dilution | Immunogen | Producer Cat # RRID | Conjugation |
|---|---|---|---|---|---|
| **Goat anti-mouse IgG (H+L)** | Goat polyclonal IgG | 1:400 | Gamma Immunoglobins Heavy and Light chains | Thermo Fisher Scientific, Cat# A32723, RRID: AB_2633275 | Alexa Fluor® 488 |
| **Goat anti-mouse IgG (H+L)** | Goat polyclonal IgG | 1:400 | Not available | Abcam, Cat# ab150118, RRID: AB_2714033 | Alexa Fluor® 555 |

| Secondary antibody | Host and type | Dilution | Immunogen | Producer Cat # RRID | Conjugation |
|---|---|---|---|---|---|
| **Goat anti-mouse IgG1** | Goat polyclonal IgG1 | 1:300 | IgG gamma 1 | Thermo Fisher Scientific, Cat# A21121, RRID: AB_2535764 | Alexa Fluor® 488 |
| **Goat anti-mouse IgG2a** | Goat polyclonal IgG2a | 1:300 | Mouse IgG2a | Thermo Fisher Scientific, Cat# A21137, RRID: AB_2535776 | Alexa Fluor® 555 |
| **Goat anti-rabbit IgG (H+L)** | Goat polyclonal IgG | 1:400 | Gamma Immunoglobins Heavy and Light chains | Thermo Fisher Scientific, Cat# A32731, RRID: AB_2633280 | Alexa Fluor® 488 |
| **Goat anti-rabbit IgG (H+L)** | Goat polyclonal IgG | 1:400 | Gamma Immunoglobins Heavy and Light chains | Thermo Fisher Scientific, Cat# A32732, RRID: AB_2633281 | Alexa Fluor® 555 |
| **Goat anti-rabbit IgG (H+L)** | Goat polyclonal IgG | 1:300 | Not available | Abcam, Cat# ab150083, RRID: AB_2714032 | Alexa Fluor® 647 |

- For the SOX2 marker, the display color was changed from green to magenta for better visibility.
